# NanoLocz: Image analysis platform for AFM, high-speed AFM and localization AFM

**DOI:** 10.1101/2023.11.23.568405

**Authors:** George R Heath, Emily Micklethwaite, Tabitha Storer

## Abstract

NanoLocz is an open-source computer program designed for high-throughput automatic processing and single-particle analysis of Atomic Force Microscopy (AFM) image data. High-Speed AFM and Localization AFM (LAFM) enable the study of single molecules with increasingly higher spatiotemporal resolution. However, efficient and rapid analysis of the images and movies produced by these techniques can be challenging, often requiring the use of multiple image processing software applications and scripts. Here, we introduce NanoLocz, an AFM and high-speed AFM analysis program that facilitates various single-particle analysis workflows through a simple, interactive interface. Workflows include but are not limited to: single-particle tracking, single-particle topographic feature analysis, single-molecule LAFM, time-resolved LAFM, and simulation LAFM. The source code and installation instructions for NanoLocz are available at https://github.com/George-R-Heath/NanoLocz.

## Introduction

Open-source tools such as RELION^1,2^ and numerous super-resolution microscopy software packages^3–8^ have played a crucial role in the ‘resolution revolutions’ of cryogenic electron microscopy (Cryo-EM) and super-resolution fluorescence microscopy (such as PALM, STORM and DNA-PAINT), respectively. The open-source and collaborative nature of these data analysis packages has fostered a vibrant community of researchers, democratizing access to advanced imaging capabilities and driving significant progress in understanding biological structures and processes. AFM and High-Speed AFM (HS-AFM) can provide video rate 3D visualization of surfaces at nanometer resolution across a range of length scales from whole cells^9^ down to single molecules.^10^ These unique capabilities enable insight into structural biology questions under various physiological conditions including protein-protein interactions^11^ and *in situ* dynamics of single molecules in response to force^12^, light^13^, ligands^14^ and drugs^15^. Furthermore, the application of single molecule localization microscopy analysis concepts to AFM data has led to the development of Localization AFM (LAFM)^16^ which enables 4Å resolution to be achieved on single biomolecules. With the significant advancements in AFM methods and hardware, the need for accessible software to harness the full potential of AFM data becomes ever more important.

The wide range of proprietary file formats provided by AFM companies (which additionally change over time) makes it particularly difficult to access, share and compare raw AFM data and apply new analysis methods. Current open-source software such as Gwyddion^17^ and WSxM^18^ enable a vast range of AFM image formats to be opened, catering to general scanning probe microscopy users but are focused on individual images. AFM image analysis tools aimed at biomolecular analysis such as TopoStats^19^ and trace_y^20^ concentrate on tracing analysis, proving powerful for the analysis of DNA and filament topology respectively. Additionally, to aid interpretation and analysis of biological structures in experimental data, tools such as the BioAFMviewer have been developed to simulate and fit AFM topographies from and to PDB structures.^21,22^ Despite these significant advancements, there remains a critical need for user-friendly AFM and HS-AFM post-processing and analysis tools that enable users of different instruments and expertise to browse, interact and analyze raw or processed AFM data, especially HS-AFM data, with high-throughput.

Here we present NanoLocz, an open-source software designed for AFM, HS-AFM and LAFM data analysis with an interactive graphical user interface. We show and explain the fundamental and new capabilities of NanoLocz which provide users with an AFM specific tool for efficient and in-depth analysis of AFM image data in diverse contexts.

## NanoLocz Workflow

Starting from a raw data AFM topography image or movie, NanoLocz is designed to have multiple possible workflows and branch points depending on the desired analysis (**Fig. 1a**). All workflows can be performed in a user-friendly graphical user interface (GUI) (**Fig. 1b**) in a semi-modular way categorized as follows:

**Fig 1.**
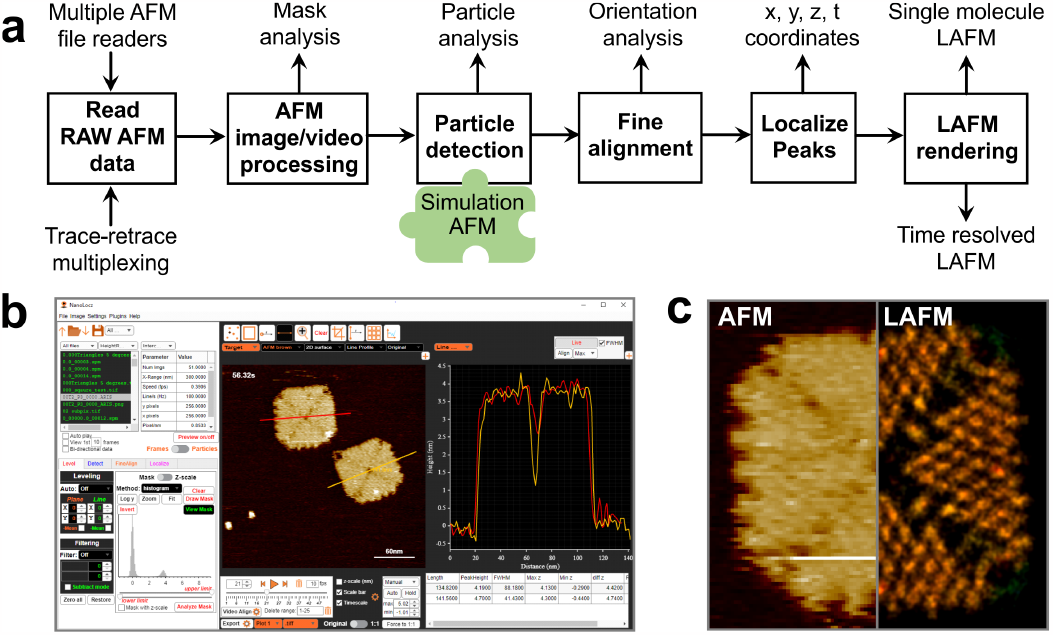
NanoLocz workflow. **a)** Overview of NanoLocz functionality and workflow demonstrated using DNA origami tiles. **b)** Graphical user interface (GUI) of NanoLocz showcasing multiple line profile analysis (red and yellow lines and traces). **c)** Comparison of a single AFM image (left) and a NanoLocz LAFM map (right) of a DNA origami tile.

### 1. Reading AFM data

Firstly, to read in raw AFM image or HS-AFM video files, multiple file formats readers are readily integrated in the software allowing topography trace, retrace scans or other data channels to be processed. The workflow of NanoLocz focuses on treating AFM data in video format, *i*.*e*., a stack of images, whilst allowing seamless integration with various AFM and HS-AFM data formats, making it suitable for data obtained from different experimental setups. See **Table 1** for the full list of accepted file formats, access to different data channels and file organization required for video format. Raw or processed AFM file formats are automatically recognized and imported as either single images or movies along with metadata of each frame. Of course, raw data is preferred as source file, as pre-processed data might already have been filtered and suffered information loss. To increase the amount of data included in the AFM analysis, trace and retrace movie frames can be loaded and intercalated in this initial step before image processing (see also **Fig. 2**). Trace-retrace intercalated movies typically require frame alignment to account for substantial offsets in the x direction. Alternatively, data which has been acquired in bi-directional scanning modes that combine trace and retrace signals in one image can be read. Bi-directional data capture can double the imaging frame rate, however it uses computational real-time data processing to align individual scan lines to build an image and is prone to contain errors that will propagate in the further data analysis particularly at higher scan rates or with higher levels of parachuting at low forces. If such data is used, NanoLocz has options for automatic correction algorithms using either line correlation or peak correlation methods.

**Table 1.**
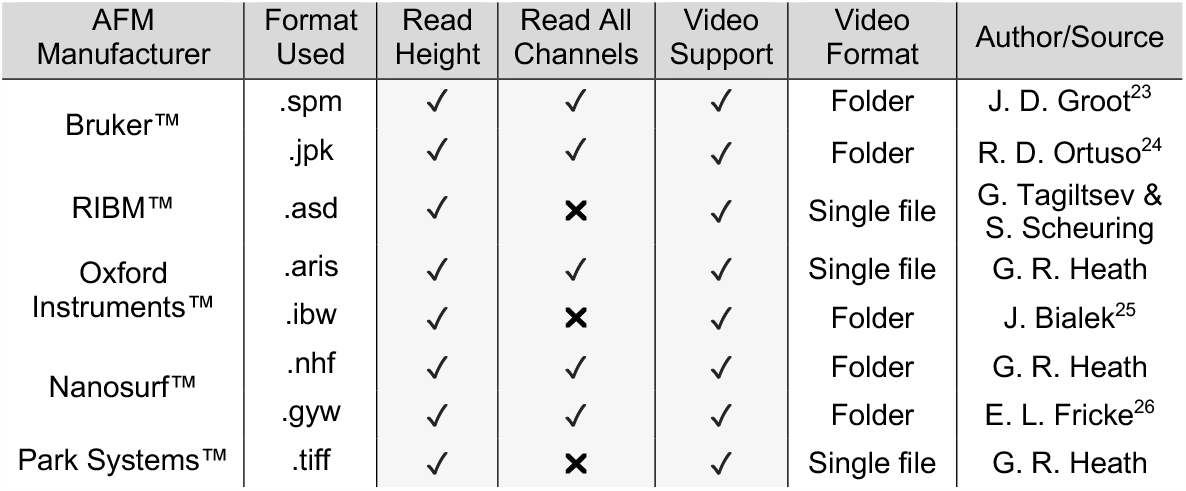
Current AFM manufacturer file formats read by NanoLocz including access to different data channels and support for video processing.

| AFM Manufacturer | Format Used | Read Height | Read All Channels | Video Support | Video Format | Author/Source |
| --- | --- | --- | --- | --- | --- | --- |
| Bruker™ | .spm | ✓ | ✓ | ✓ | Folder | J. D. Groot<sup>23</sup> |
|  | .jpk | ✓ | ✓ | ✓ | Folder | R. D. Ortuso<sup>24</sup> |
| RIBM™ | .asd | ✓ | ✗ | ✓ | Single file | G. Tagiltsev & S. Scheuring |
| Oxford Instruments™ | .aris | ✓ | ✓ | ✓ | Single file | G. R. Heath |
|  | .ibw | ✓ | ✗ | ✓ | Folder | J. Bialek<sup>25</sup> |
| Nanosurf™ | .nhf | ✓ | ✓ | ✓ | Folder | G. R. Heath |
|  | .gyw | ✓ | ✓ | ✓ | Folder | E. L. Fricke<sup>26</sup> |
| Park Systems™ | .tiff | ✓ | ✗ | ✓ | Single file | G. R. Heath |

**Fig 2.**
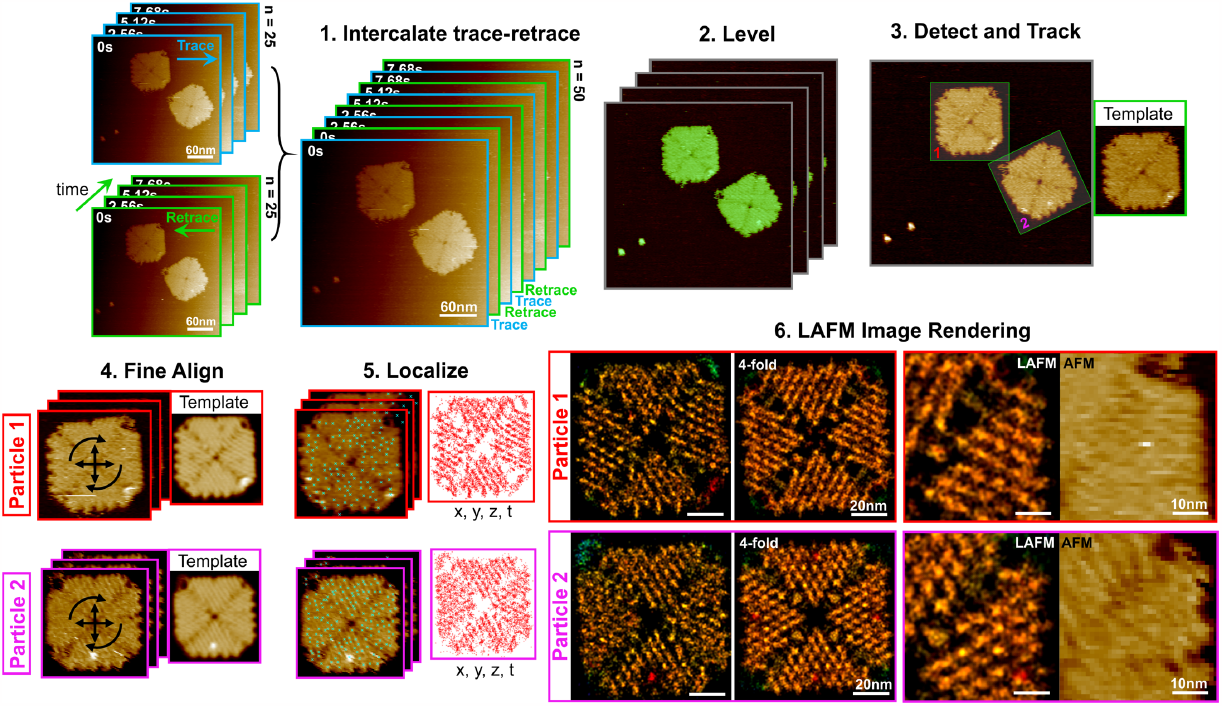
Example single particle LAFM workflow. 1. Loading of raw data and intercalation of trace and retrace frames. 2. Iterative levelling of the movie frames with height thresholding. 3. Detection of 4FS DNA origami tiles using a user selected reference image as a template (a single 4FS Origami tile image, insert on the right). 4. Iterative fine alignment of each individual particle using an averaged image as template updated after each iteration. 5. Localization of local height fluctuations to generate a 3D point cloud of localizations in x, y, z and time. 6. Rendered localization AFM images without (left) and with (right) 4-fold rotational symmetry averaging. Right: zoomed comparison between symmetrized LAFM map and single particle AFM images.

### 2. Image Processing

Raw AFM images typically require background and line leveling to correct for sample tilt and height variations between scan lines. This process often requires multiple iterations to fit data with higher or lower height features excluded to avoid skewing the leveling. NanoLocz increases throughput allowing automatic image leveling with pre-set routines and real-time previews during movie processing, eliminating the need for slow batch processing and leveling errors. Leveling can also be performed manually. The manual and automatic thresholding algorithms account for height throughout a movie or set of images to avoid changes in baseline and improve accuracy. At this step in the image processing pipeline one can perform standard AFM analysis such as threshold area/height analysis and line profiling, or advanced further towards single particle analysis.

### 3. Particle Detection

Depending on the sample and resolution achieved in AFM images, single particles may appear as featureless amorphous blobs or display visible substructure. Single particle detection and analysis in NanoLocz can be performed using either local maxima (for featureless objects) or using cross-correlation searches with a reference image (for particles with visible substructure). Using the local maxima method automatically provides an output of statistics on particle height, location and width using automated height profiling. Alternatively for applications that require analysis of the particle substructure such as LAFM, reference based single particle detection can be performed. NanoLocz implements algorithms that enable detection using a reference region in the image, *e*.*g*., a typical particle, selected by the user, or using a simulated reference image using a procedure that simulates an AFM topography from a 3D coordinates (for example a protein structure from the Protein Data Bank, *i*.*e*., PDB file). To develop a multi-rotation search algorithm, we combine correlation coefficient maps from the different reference images taking their maxima at each x, y position, the local maxima in the combined map is then matched back to the rotation parameters used to create the template for detection. The user interface facilitates interactive tuning of particle detections by employing a slider to visualize in real-time the effect of changing the minimum correlation coefficient and/or height thresholds, enabling rapid visual feedback for the inclusion or exclusion of detected particles. If required, particle detection can then be refined using an averaged image of the particles detected in this first round of cross-correlation searching. Once detected, single particle tracking can be run to identify and follow particles over time. At this stage statistics of particle angle, height, spatial distribution and movements over time can also be output.

### 4. Fine Alignment

Once positions have been detected particles can be rotationally and translationally aligned further. Translational and rotational alignment to the reference image is performed in multiple iterations in which the detection positions and reference are refined in each iteration. To prevent data loss, each alignment iteration updates the detection’s x, y coordinates and angle only. This approach allows for a one-time particle extraction from the full image, avoiding losses that may occur due to multiple pixel interpolations in the sequential iterations. If tracking has been performed, alignments can be made to either all particles or a single particle over time.

### 5. Localization AFM

Once the particle stack (either many individual molecules, or many observations of one or several molecules over time) is aligned, it can be utilized for LAFM. The first step of LAFM is to find local height maxima caused by height fluctuations whilst avoiding background noise. To achieve this, the local maxima detected in response to non-destructive image filters (without modification of the image data) can be previewed, along with adjusting thresholds for height and peak prominence. A typical recommended filter strength is a Gaussian filter with a strength of at least 0.6. To allow refinement of the peak detection positions to subpixel localizations, bicubic interpolation is applied to a 5x5 subset of pixels surrounding the initial local maxima, expanding it to 50x50 pixels. The maxima of these interpolated subsections are then used as the final localizations. For each localization, information about on x, y, z, time, and peak prominence is stored. The LAFM image can then be rendered using these localizations on an expanded pixel grid with points rendered using 2D Gaussian profiles and 2D false-color scale options which combine height and localization probability. For molecules with symmetry, a symmetrized LAFM map can be generated with automatic centering based on the alignment of the rotated localization point cloud. At this stage LAFM images can be rendered to produce time resolved LAFM movies and/or single molecule LAFM images. **Fig. 1c** demonstrates a comparison between a single AFM image and the LAFM map obtained through NanoLocz processing.

### 6. Data Export

Processed images and movies in NanoLocz can be saved and opened later or exported to a range of file formats. Exporting as .tiff, .csv, .txt, or .xls enables export without loss of image information whereas export as .gif, .avi, .png, .jpeg or .pdf gives movies/images at presentation or publication quality with automatic scale bars and timestamps. All other data obtained through NanoLocz analysis, such as height and width distributions, particle orientations, tracking coordinates and localizations can also be exported and plotted within the application.

## Example Workflow: Single Particle LAFM

To demonstrate the capabilities of NanoLocz, **Fig. 2** shows an example workflow to obtain single particle LAFM images of two different copies of the same 4FS DNA origami tile design.^27,28^ The raw data contains 25 images in which the two DNA origami tiles are visible. Typically, LAFM imaging requires at least 100 images of the same or different particles to get a reasonable sampling of localizations.^16^ To increase the number of images for LAFM analysis, trace-retrace frame intercalation was used to give 50 unique frames and therefore 50 images for each particle. Given that the 4FS Origami tile has 4-fold symmetry, we actually pool the information of 100 (trace only) or 200 (trace and retrace) redundant structural elements in our LAFM maps. After leveling the movie, DNA origami tiles were detected using a reference region from the leveled images with rotational freedom enabled. Particles were then tracked allowing separation of particle 1 and 2 for image fine alignment using single particle averages to iteratively refine alignment further. Each particle can then be treated separately in the localization analysis to generate independent localization point clouds and subsequent LAFM images. The LAFM images show positions of defects unique to each origami. 4-fold symmetrization of the two single particle LAFM images shows the matching structural features of the DNA origami organization at high-resolution.

Comparison of the LAFM images to the raw AFM data images highlight the significant improvement in resolution and contrast gained through the LAFM processing.

## Example Workflow: Simulation AFM Reference and Time-Resolved LAFM

Integration of AFM topography simulation software into NanoLocz enables the semi-automatic generation and import of simulated AFM surfaces, using pixel sampling matching the experimental data, from protein structures in repositories such as the Protein Data Bank. Once imported these simulated surfaces can be used as a reference image for particle detection, to find protein location and orientation or to test LAFM on a simulated protein surface. In **Fig. 3a** a CLC-ec1 antiporter structure (PDB: OST1^29^) is used to produce a simulated molecular topography. In brief, AFM surface topography prediction of the extracellular CLC-ec1 face is based on scanning *in silico* an estimated tip radius over the PDB structure using the identical pixel sampling as in the experimental data. This simulated surface is then used as a template for cross-correlation searches and particle detection in the HS-AFM experimental data. The simulated template is also used as reference for fine rotational and translational alignment. The data obtained from the fine alignment such as x, y position, rotation angle and normalization cross-correlation value (with the simulated surface) gives information about the positional dynamics and conformational variations (or image quality) over time (**Fig. 3b**). Such analysis, potentially using several simulation topographies of the protein in different states, could also be used to detect conformational changes over time. When performing localization analysis NanoLocz stores the time point of each localization which can then be used to produce LAFM movies when rendering at user defined numbers of particles time (**Fig. 3b**). Whilst the CLC-ec1 molecule is not undergoing significant conformational changes on the 18s timescale, such a time resolved LAFM workflow can be applied to HS-AFM which contains multiple copies of the same particle in each frame to increase the effective LAFM time resolution.

**Fig 3.**
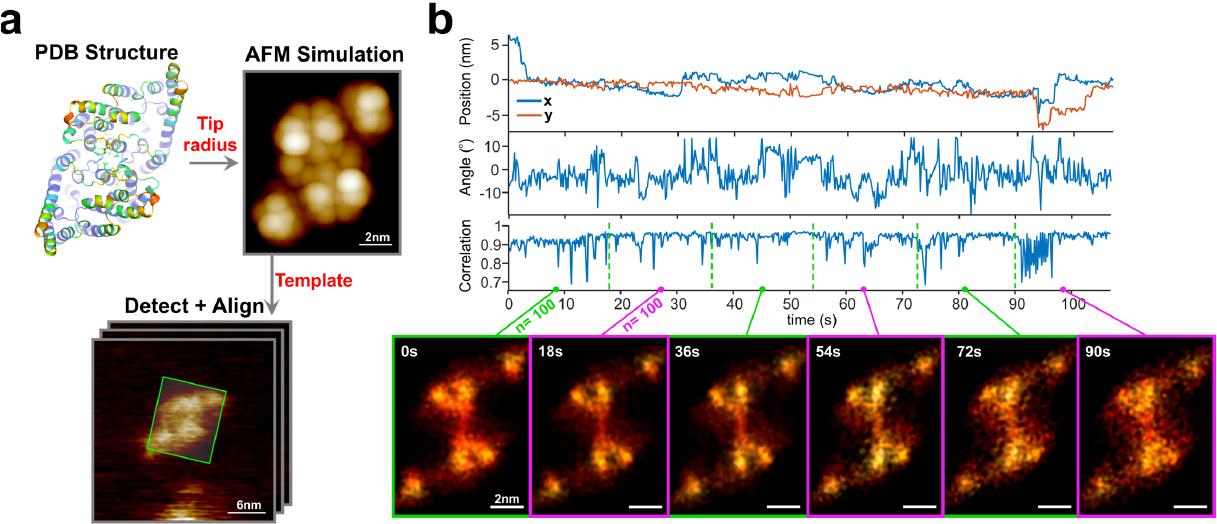
Example Simulation AFM and Time-Resolved LAFM Workflow. **a**) Demonstration of the use of an AFM simulation image, the extracellular face of CLC-ec1 (PDB 1OTS^29^), as reference, imported into NanoLocz for particle detection and fine alignment. **b**) x, y detection position, angle and normalized cross-correlation of the HS-AFM experimental particle over time compared to the simulated structure reference. Time resolved LAFM maps of the CLC at 18s intervals using 100 independent frames per LAFM map. LAFM maps have 2-fold symmetry applied.

## Conclusions

NanoLocz’s advanced capabilities and user-friendly interface make it a valuable tool for AFM image analysis. By streamlining and integrating AFM, HS-AFM and LAFM data processing workflows, NanoLocz significantly reduces manual efforts which should lead to increased productivity, more access to single molecule analysis methods and help pave the way for future advances in AFM analysis. The ongoing development and future enhancements to NanoLocz including python versions aim to further support new improvement in AFM data analysis capabilities. By encouraging the development of novel analysis algorithms through plugin integration, NanoLocz aims to create a collaborative environment for further advancements in AFM research. Such an open-source non-proprietary software package used by the AFM community at large will automatically lead to common standards and enhance data reproducibility and exchange.

## Code Availability

The source-code, standalone versions for PC or Mac, MATLAB app, test data and installation instructions for updated versions of NanoLocz and can all be found at https://github.com/George-R-Heath/NanoLocz.

## Acknowledgements

We thank Abeer Alshammari and Maya Tekchandani for extensive testing of the software, Grigory Tagiltsev and Simon Scheuring for providing .asd file opening algorithms, Hector Corte for assisting with .nhf and .gwy file opening algorithms. Christoph Walti and Ilaria Sandeifor for providing 4FS DNA origami tiles. We gratefully acknowledge funding from the Engineering and Physical Science Research Council in an EPSRC Open fellowship (EP/W034735/1) to G. R. Heath.

## Notes

### Competing Interest Statement

The authors have declared no competing interest.

